# Oral administration of 4-methylumbelliferone reduces glial scar and promotes anatomical plasticity

**DOI:** 10.1101/2023.02.02.526565

**Authors:** Kateřina Štepánková, Milada Chudíčková, Zuzana Šimková, Noelia Martinez-Varea, Šárka Kubinová, Lucia Machová Urdzíková, Pavla Jendelová, Jessica C F Kwok

## Abstract

Following a spinal cord injury (SCI), chondroitin sulfate proteoglycans (CSPGs) are up-regulated at the glial scar inhibiting neuroregeneration. Under normal physiological condition, CSPGs interact with hyaluronan (HA) and other extracellular matrix on neuronal surface forming a macromolecular structure called perineuronal nets (PNNs) which regulate neuroplasticity. 4-methylumbelliferone (4-MU) has been used previously to down-regulate HA synthesis but not been tested in SCI. In this study, we have evaluated the effect of 4-MU, an inhibitor of HA, in a chronic contusion model of SCI in rats. At a dose of 1.2 g/kg/day of 4-MU, we observed not only the reduction of HA in the uninjured spinal cords after 60 days of 4-MU administration, but also a down-regulation of CS glycosaminoglycans (CS-GAGs). In order to assess the effect of 4-MU in chronic SCI, rats with T8 spinal contusion injury were fed with 4-MU or placebo for 8 weeks in combination with daily treadmill rehabilitation for 16 weeks to promote neuroplasticity. 4-MU treatment promoted significant sprouting of 5-hydroxytryptamine (5-HT) positive fibres into ventral horns and reduced the HA synthesis by astrocytes around the lesion site. While 4-MU reduced astrogliosis in chronic stage of SCI, the current dose was not sufficient to down-regulate the increased production of CS-GAGs or behavioural performance. Together, these data suggest that oral treatment with 4-MU is able to induce anatomical plasticity but further adjustment on the dosage will be required to benefit functional recovery after SCI.

## INTRODUCTION

Spinal cord injury (SCI) is a damage to the spinal cord that causes partial or complete loss of control of locomotor and sensory functions (Keough *et al*. 2016). SCI itself is a dynamic process starting immediately after the injury when tissue damage is continued with haemorrhage, inflammation, as well as oedema. The acute phase includes initial trauma followed by spinal cord ischemia, cell excitotoxicity, ion dysregulation, and free radical-mediated peroxidation (Pinchi *et al*. 2019). These processes initiate a complex secondary injury cascade leading to changes in structural architecture of the spinal cord characterised by the chronic phase. Chronic phase begins 6 months after SCI in human and continues throughout the lifetime of the patient. It is characterized by the stabilization of the lesion including scar formation accompanied by alterations in neural circuitries (Oyinbo 2011). As there is no successful regenerative treatment available, most patients remain in the chronic state for the rest of their life.

Adult central nervous system (CNS) has relatively poor regeneration capacity caused by both the extrinsic inhibitory environment after injury (such as glial scar and myelin-associated inhibitors) and the intrinsic poor regeneration ability of the neurones themselves (Kaplan *et al*. 2015; Vogelaar 2016). While these extrinsic and intrinsic factors co-ordinate to maintain the stability of the neuronal networks in a healthy state, they rapidly become the major obstruction to regeneration post-injury (Nagappan *et al*. 2020). Perineuronal nets (PNNs) are extracellular matrix (ECM) structures enwrapping a sub-population of neurons in the CNS and play a crucial role in plasticity regulation during postnatal development and post traumatic regeneration (Carulli and Verhaagen 2021; van ‘t Spijker and Kwok 2017). PNNs form at the end of the critical period and terminate developmental plasticity. PNNs mature by early adulthood (Pizzorusso *et al*.2002) and stabilize synaptic contacts and are dynamically maintained during the lifespan (Pizzorusso *et al*.2002; Carulli *et al*. 2010; Tsien 2013). In spinal cord, PNNs mainly surround motoneurons in the ventral horns (Takahashi-Iwanaga *et al*. 1998; Vitellaro-Zuccarello *et al*. 2007; Galtrey *et al*. 2008) that are directly responsible for the motor activity (Stifani 2014). PNNs are composed of a multitude of neural ECM components where chondroitin sulphate proteoglycans (CSPGs) and hyaluronan (HA) are the major constituents (Giamanco *et al*. 2010; Irvine and Kwok 2018; van ‘t Spijker and Kwok 2017). Lecticans, a sub-family of HA binding CSPGs, bind to the long HA chains forming a meshwork on neuronal surface and that this interaction is stabilised by the HA and proteoglycan link proteins (Haplns) (Djerbal *et al*. 2017).

Degradation of PNNs reactivate juvenile-like states of plasticity which enable axon sprouting and regeneration of function by synaptogenesis and experience-dependent synaptic plasticity after SCI (Sorg *et al*. 2016). Enzymatic removal of PNNs through chondroitinase ABC (ChABC), that digests chondroitin sulphate glycosaminoglycan (CS-GAG) chains on CSPGs, has shown to reopen a critical window for plasticity enhancement and promote functional recovery in multiple SCI models (Bradbury *et al*. 2002; García-Alías *et al*. 2009). Moreover, when ChABC is combined with rehabilitation, the effect of ChABC treatment is enhanced as compare to non-rehabilitating animals (Smith *et al*. 2015; Al’joboori *et al*. 2020). CSPG synthesis inhibitors, such as fluorosamine and its analogues, have also been shown to reduce CSPG level and accelerate remyelination following focal demyelination in mice (Keough *et al*. 2016; Stephenson *et al*.2019). These results suggest that CSPG synthesis inhibitors are possible approaches to enable axonal regeneration after CNS injury.

We have recently reported the use of a small molecule 4-methylumbelliferone (4-MU) to down-regulate PNNs and re-activate neuroplasticity for memory enhancement (Dubisova *et al*. 2022). 4-MU is a known inhibitor to HA synthesis (Nagy *et al*. 2015; Nagy *et al*. 2019). Previous work has shown that 4-MU inhibits the production of UDP-glucuronic acid (UDP-GlcA) (Galgoczi *et al*. 2020), a key substrate for HA production, and the expression of hyaluronan synthases (HASs), UDP-glucose-pyrophosphorylase (an enzyme for the biosynthesis of UDP-glucose) and UDP-glucose-dehydrogenase (an enzyme that converts UDP-glucose into UDP-glucuronic acid (Kakizaki *et al*. 2004; Kultti *et al*. 2009). Interestingly, UDP-GlcA is also a substrate for the synthesis of CS, as well as some other GAGs including dermatan and heparan sulphates. The effect of 4-MU in general GAG synthesis has yet to be clarified.

Our previous observation of PNN down-regulation by 4-MU prompts us to ask the question if 4-MU would be effective in down-regulating the inhibitory ECM (i.e. ECM molecules, such as proteoglycans, which are up-regulated after injury and restricting axonal growth) in the glial scar after SCI. Here, we aimed to evaluate the biochemical and histological changes in the injured spinal cord after long-term 4-MU treatment in rats and to evaluate the effect of this molecule on anatomical changes in the spinal cord after chronic SCI. Histological staining of PNNs using *Wisteria floribunda agglutinin* (WFA) and anti-aggrecan antibody (ACAN, a CSPG) shows PNN down-regulation in 4-MU-treated group, and that the staining recovers in wash-out group. We also observed that the long-term 4-MU treatment leads to a significant reduction in the glial scar and promotes the sprouting of serotonergic fibres above and below the lesion.

## METHODS

### Experimental animals

55 female Wistar RjHan:WI rats (8-week-old, 250 g–300 g; 2020-2021; CS 4105 Le Genest Saint Isle; Saint Berthevin Cedex 53941 France) were used in total. 40 animals were used for the histochemical and biochemical assessment of the 1.2 g/kg/day 4-MU dose effect in 4-MU-fed/non-fed combined with/without rehabilitation groups. These animals were divided into 5 groups (8 rats per group) using simple randomization – placebo, placebo + training/rehabilitation, 4-MU, 4-MU + rehabilitation, and 4-MU with planned 2 months post-feeding period (wash-out group). Based on our previous experiments, we are expecting an effect size of ≥1.7. A power calculation of α=0.05, number of groups = 5, a total sample size of 8 is required for each experimental group.

Hereafter, the histochemical and biochemical assessment of the treatment in uninjured animals was done, 18 animals underwent the spinal cord contusion, 3 animals with BBB score less than 8, one week after the injury were excluded. The feeding has begun 6 weeks after SCI when the chronic phase was fully established. Half of the animals received chocolate-flavoured chow containing 4-MU (1.2 g/kg/day) (treated group) and the second half just chocolate-flavoured chow without any treatment (placebo group). Placebo and treated groups received daily physical rehabilitation on treadmill. Feeding stopped after 8 weeks, but daily extensive rehabilitation continued for 2 more months.

Rats were housed by two in cages with 12 hours light/dark with standard conditions (in temperature (22 ± 2°C) and humidity (50% ± 5%)). Rats had free access to water and food *ad libitum*. All experiments were performed between 08:30 to 19:00 local time.

### Animal surgeries

Animals received a severe thoracic spinal cord contusion using a commercially available Infinite Horizon (IH) spinal cord injury device (IH-0400 Spinal Cord Impactor device; Precision Systems and Instrumentation, Lexington, KY, USA). All procedures were approved by the ethical committee of the Institute of Experimental medicine of ASCR and performed in accordance with Law No. 77/2004 of the Czech Republic (Ethics approval number: 13/2020). Power calculation, based on previous studies, was performed prior to the experiment to estimate the number of animals needed. All work was performed according European Commission Directive 2010/63/EU and all efforts were made to minimize pain and suffering. Rats were anesthetized with 5% (v/v) isoflurane and maintained at 1.8-2.2% during the surgery, in 0.3 L/min oxygen and 0.6 L/min air. Animals were injected subcutaneously with buprenorphine (Vetergesic_^®^_ Multidose, 0.2 mg/kg body weight). Using sterile procedures, a laminectomy was performed at the Th8/Th9 level with the vertebral column being stabilized with Adson tissue forceps at T7 and T10. The animals received 200 kilodynes (kdyn) moderate contusion injury. Manual expression of bladders was performed in the first two weeks after injury, and the rats were checked daily. Rats were given treatment of analgesics if signs of inflammation were observed. Animals were given a random number after the surgeries. The chow and the placebo was similarly given a random number, and fed to the animals according to the number. Experimenters were blind to the treatment group during behavioural analysis. Identity of the animals and their treatment group were only revealed after assessment were performed.

At the end of the experiments, half of rats from each group was intraperitoneally anesthetized with a lethal dose of ketamine (100 mg/kg) and xylazine (20 mg/kg), perfused intracardially with 4% (w/v) paraformaldehyde (PFA) in 1X-PBS and post-fixed in the same solution for 24 hours. Spinal cords were dissected before storing in 30% (w/v) sucrose (Milipore, cat. no. 107651) in 1X PBS. The second half of the rats were sacrificed with a lethal dose of ketamine (100 mg/kg) and xylazine (20 mg/kg) before the spinal cords were dissected and froze on dry ice before storage at −80°C for subsequent qPCR and GAGs extraction. The anaesthesia used have previously been shown not to interfere with animal behaviour.

### 4-MU treatment

2.5% (w/w) 4-MU was mixed and prepared into rat chocolate-flavoured chow (Sniff GmbH, Germany). This percentage would allow the delivery of 4-MU at 1.2 g/kg/day to the rats when chow was consumed *ad libitum*, an amount assessed from our pilot experiment. Rats were fed with the 4-MU chow at 10 weeks of age (non-SCI cohort) or 6 weeks after the SCI (SCI-cohort). Rats were maintained on 4-MU diet for 8 weeks. The food intake was measured weekly to check for consumption (Figure 1).

**Figure 1.**
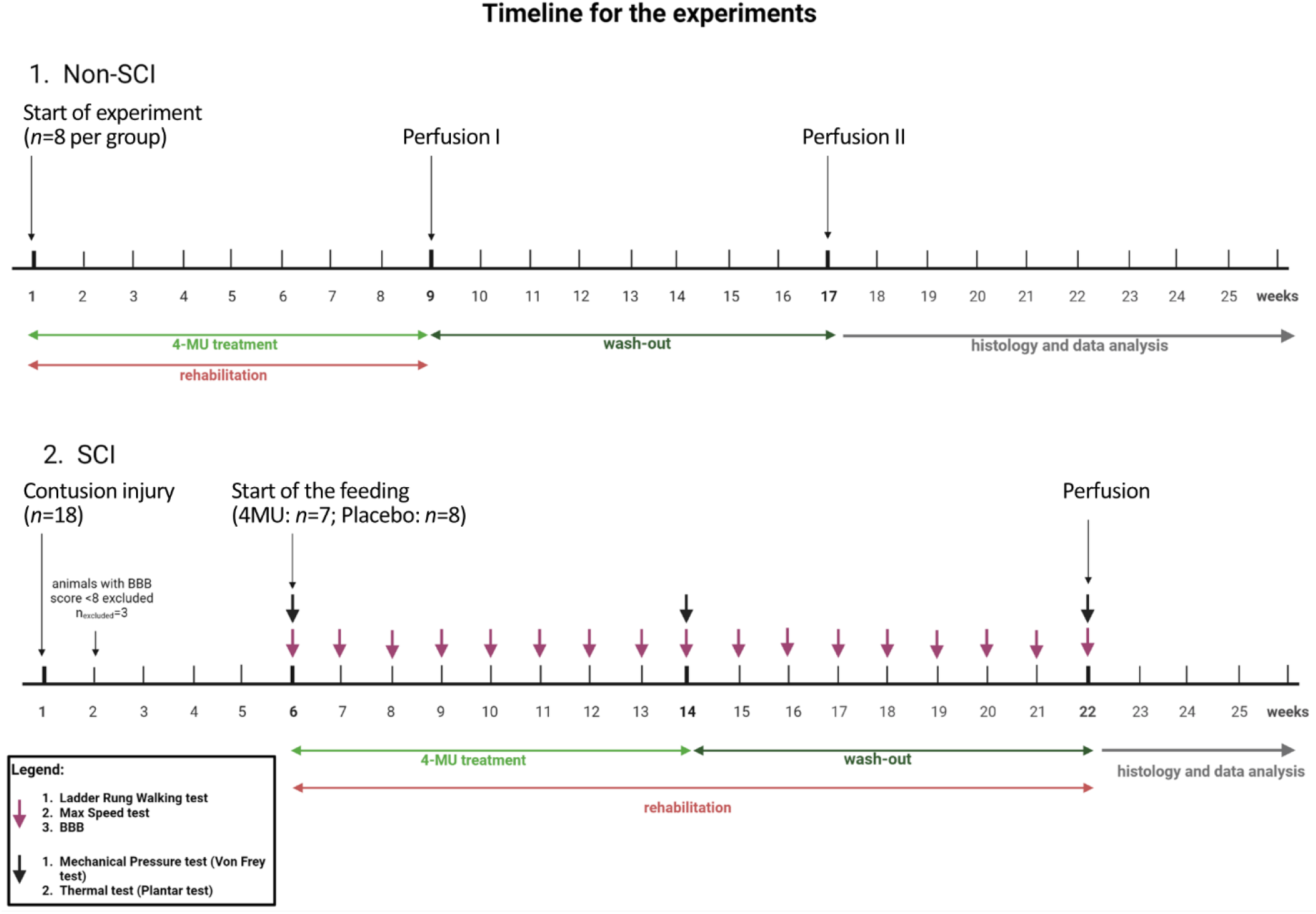
Schematic illustration of the experimental timeline. The timeline of the experiments was created with BioRender.com.

### Histological and histochemical analysis

**Two cm long spinal cords, with the Th8 level in the middle, were embedded in O.C.T. compound (VWR) and sectioned in frozen block of tissue before being sectioned into 40 μm thick slices for immunohistological** analysis. The sections were made in series of 1 mm, compared to the anatomical atlas, and thus the spinal cord segments were identified.

40 μm sections (free-floating for uninjured spinal cords, and mounted on glass slides for injured spinal cords) were permeabilizated with 0.5% (v/v) Triton X-100 in 1X PBS for 20 min and then endogenous biotin was blocked using Avidin/Biotin blocking kit (Abcam; cat. no. ab64212) to reduce non-specific background. Tissue was then blocked in ChemiBLOCKER (1:10; Millipore cat. no. 2170), 0.3 M glycine, 0.2% (v/v) Triton X-100 in 1X PBS for 2 hrs. The sections were then incubated with labelling agents including biotinylated *Wisteria floribunda* agglutinin (1:150, 24 hrs), biotinylated hyaluronan binding protein (HABP) (1:150, 24 hrs) and/or primary antibodies: anti-choline acetyltransferase (ChAT)(1:600, 72 hrs); anti-ACAN (1:150, 24 hrs), anti-GFAP conjugated to Cy3 (1:800, 24 hrs), anti-5HT (1:100; 48 hrs) or anti-synapsin (1:100; 48 hrs) as shown in Table 1. After washing, the sections were labelled with fluorescent-conjugated secondary antibodies (1:300; 2 hrs; room temperature (RT) (table 1). Staining was imaged with a LEICA CTR 6500 microscope with FAXS 4.2.6245.1020 (TissueGnostics, Vienna, AT) software. Images were evaluated for the total number of cells in ventral horns with WFA-positive signal, ACAN-positive signal and number of co-localised cells using ImageJ^™^ (NIH, Bethesda, MD, USA). The HABP intensity was analysed with HistoQuest 4.0.4.0154 (TissueGnostics) software. 5HT receptor and synapsin staining was imaged with confocal microscope Olympus FV10i (Olympus Life Science, Waltham, MA, USA). The intensity of 5HT and synapsin was measured in the ventral horn by ImageJ^™^ and compared between 4-MU-treated and placebo group. All images for comparison were taken using identical settings and under the same staining conditions.

**Table 1.** A table showing the primary antibodies and fluorescent-conjugated secondary antibodies used for the experiments.

| 1° Antibody/labelling agent | Epitope | Host | Company |
| --- | --- | --- | --- |
| Biotinylated *Wisteria floribunda* agglutinin (WFA) | Chondroitin sulfate in the PNNs | N/A | Sigma-Aldrich cat. no. L1516 |
| Biotinylated Hyaluronan Binding protein (b-HABP) | Hyaluronan | N/A | Amsbio cat. no. AMS.HKD-BC41 |
| Anti-ACAN | Aggrecan | Rabbit polyclonal | Sigma-Aldrich cat. no. AB1031 |
| Anti-GFAP-Cy3 | Glial fibrillary acidic protein | Mouse monoclonal IgG | Sigma-Aldrich cat. no. C9205 |
| Anti-ChAT | Choline acetyltransferase | Mouse monoclonal IgG | Invitrogen cat. no. MA5-31383 |
| Anti-5HT2C | 5-hydroxytryptamine receptor | Rabbit monoclonal | Novus Biologicals cat. no. NBP2-67100 |
| Anti-synapsin | Synapsin | Rabbit polyclonal | Novus Biologicals cat. no. NB300-104 |
| 2° Antibody/labelling agent |  | Company |  |
| Streptavidin-Alexa fluor 488 |  | Invitrogen cat. no. S32354 |  |
| Goat anti rabbit Alexa fluor 594 |  | Invitrogen cat. no. A11012 |  |
| Goat anti mouse Alexa fluor 405 |  | Invitrogen cat. no. A48255 |  |
| Goat anti rabbit Alexa fluor 594 |  | Invitrogen cat. no. A11012 |  |
| Goat anti rabbit Alexa fluor 594 |  | Invitrogen cat. no. A11012 |  |

### GAGs extraction and analysis

Frozen tissues were first weighted before the incubation with acetone to remove the lipids. The samples were then dried and cut into small pieces before the pronase treatment (15 mg pronase per hemisphere, (Roche, cat. no. 11459643001) in 0.1 M Trizma hydrochloride (Sigma-Aldrich, cat. no. T3253), 10mM calcium acetate (Millipore, cat. no. 567418), pH 7.8. Samples were homogenized with Potter Elvehjem tissue homogenizer in the pronase solution. Residual protein fragments were precipitated with trichloroacetic acid (Sigma-Aldrich, cat. no. T6399). The supernatant, which contains the GAGs, was collected and stored on ice. The solution was then neutralised with 1 M Na_2_CO_3_ (Sigma-Aldrich, cat. no. S7795) to pH 7.0, and the GAGs were recovered by ethanol precipitation. The isolated GAGs were redissolved in 0.3 ml of deionised water. The GAG concentration was quantified using cetylpyridium chloride (CPC) turbidimetry. Standard curve was prepared from 1μg/μl of Chondroitin sulphate A (Sigma-Aldrich, cat. no. C9819) (Kwok *et al*. 2015). Briefly, the diluted sample was mixed with 0.2% (w/v) CPC and with 133 mM MgCl_2_ (Sigma-Aldrich, cat. no. M2670) in ratio 1:1. Absorbance was measured at 405 nm using plate reader spectrophotometer (FLUOstar^®^ Omega, BMG LABTECH). Each sample was carried out in three different dilutions and each dilution was carried out in duplicate.

### Quantitative real-time polymerase chain reaction

For quantitative evaluation of gene transcript levels in spinal cords treated with or without 4-MU, quantitative real-time PCR (qPCR) was used. RNA was isolated with RNeasy_©_ Lipid Tissue Mini Kit (QIAGEN, cat. no. 74804) according to the manufacturer’s protocol. Amount of isolated RNA was quantified using NanoPhotometer_©_ P330 (Implen, München, Germany). Then, TATAA GrandScript cDNA Synthesis Kit (TATAA Biocenter, Art No. AS103c) was used for reverse transcription of RNA into complementary DNA (cDNA), following the manufacturer’s protocol in the T100TM Thermal Cycler (Bio-Rad, Hercules, CA, USA). For the qPCR, TaqMan^®^ Gene Expression Assays (Life Technologies by Thermo Fisher Scientific, Waltham, MA, USA) were used for HAS1 (Rn01455687_g1), HAS2 (Rn00565774_m1), HAS3 (Rn01643950_m1) and GAPDH (Rn01775763_g1), all purchased from Applied Biosystems and used as recommended by the manufacturer. Amplification was performed on the qPCR cycler (QuantStudio^™^ 6 Flex Real-Time PCR System, Applied Biosystems^®^ by Thermo Fischer Scientific, Waltham, MA, USA). All amplifications were run under the same cycling conditions: 2 min at 50 °C, 10 min at 95 °C, followed by 40 cycles of 15 s at 95 °C and 1 min at 60 °C. Expression was calculated using the threshold cycle (Ct) value and log(2^-ΔΔCt) method and normalized to the control group. Each qPCR experiment was carried out in duplicate. Ct values of each measured condition were normalized to glyceraldehyde 3-phosphate dehydrogenase (GAPDH). Then, the 2^-ΔΔCt^ values were expressed as described by Livak and Schmittgen (Livak and Schmittgen 2001). After that, the mean of each subset was calculated for each group.

### Behavioural tests

#### Treadmill training

Treadmill training began 6 weeks after SCI and was conducted on 5 consecutive days per week for 16 weeks. The training consisted of 10 minutes run, 20 minutes break and 10 minutes run. The treadmill speed was 16-18 cm/s in the first and second weeks and 20 cm/s in the remaining weeks of training. Half of non-SCI rats were trained for 8 weeks as well, except wash-out group when the training had been prolonged for 2 months without 4-MU feeding.

#### Max Speed test

Max Speed test was performed once per week and began 6 weeks after the SCI. The treadmill speed started at 20 cm/s and was increased every 10 s for 2 cm/s. When the animal was not able to run at a given speed, the treadmill was stopped and the last value was recorded.

#### Basso, Beattie and Bresnahan (BBB)

The Basso, Beattie and Bresnahan (BBB) (Basso *et al*. 1995) open-field test was used to assess the locomotor ability of the rats. The rats were placed into the arena bordered with rectangular-shaped enclosing. The results were evaluated in the range of 0–21 point; from the complete lack of motor capability (0) to healthy rat-like locomotor ability (21). The measurements were performed 4^th^ and 7^th^ day after the SCI and then weekly for 8 weeks, starting 6 weeks after SCI.

#### Ladder Rung Walking test

For the advanced locomotor skills, the ladder rung walking test had been used. Animals were placed on a 1.2 m – long horizontal ladder, with irregularly spaced rungs. At the end of the ladder, dark box rats were familiar with was placed. The animals crossed the ladder 3 times in a row and all attempts were recorded on camera. The hind paws placement on the rungs was evaluated by using a seven-category scale (0–6 points), as previously described by Metz and Whishaw (Metz and Whishaw 2009) from all three videos. Metz and Whishaw’s scoring scale was divided into 3 categories: 0-2; 3-4; 5-6. Videos were then evaluated and the percentage of steps in each of the 3 categories was calculated. All the animals were pretrained before lesioning. The test was performed weekly for 16 weeks, starting 6 weeks after the SCI.

#### Mechanical Pressure test (Von Frey test)

The Mechanical Pressure test was used to assess mechanical allodynia after the SCI and/or during treatment. The animals were placed in an enclosure with a metal mesh bottom and habituated there for 15 minutes. Von Frey rigid tip coupled with a force transducer (IITC Life Science, California, USA) was applied with a gradual increase of pressure to the footpad of the forepaw, until the animal withdrew its paw. The maximum pressure was recorded in grams. Five trials were performed on both hind paws. The trial was terminated if the animal failed to respond within 90 g. The average was set from three values after excluding the highest and lowest measurement. The test was performed 6 weeks after the SCI (before 4-MU treatment), after 8 weeks of rehabilitation (after 4-MU treatment), and at the end of the experiment.

#### Thermal test (Plantar test)

The Thermal test was used to assess a thermal hyperalgesia after the SCI and/or during treatment. The animals were placed in an acrylic box of the standard Ugo Basile test apparatus (Ugo Basile, Comerio, Italy) and habituated there for approximately 30 minutes. A mobile infrared-emitting lamp was then placed directly under the footpad of the hind paw, always in the same position. After placing the lamp, a thermal radiant stimulus was applied. The apparatus is connected to device what automatically recorded the time (in seconds) between the outset of the stimulus and the paw-withdrawal. Five trials were performed on both hind paws. The trial was terminated if the animal failed to respond within 30 s or wet oneself. The average was set from three values after excluding the highest and lowest measurement. The test was performed 6 weeks after the SCI (before feeding), after 8 weeks of rehabilitation (after feeding), and at the end of the experiment.

### Statistical analysis

Data processing and statistical analysis were performed using GraphPad Prism (GraphPad Software). To analyse the effect of 4-MU treatment on chronic SCI in rats, different statistical tests were used. The two-way *ANOVA* followed by Sidak’s multiple comparisons test was applied for astrogliosis, HABP, WFA, ACAN, 5HT, synapsin intensity in SCI cohorts, BBB and Ladder Rung Walking test. The two-way *ANOVA* followed by Dunnett’s test multiple comparisons test was applied intensity of HABP, number of cells enwrapped by PNNs within intact spinal cords, and for the gene expression of *has* genes. Different *post-hoc* tests for two-way ANOVA were used. Dunnett’s test was used in order to test specifically the 4-MU effect and/or rehabilitation effect against a reference group (placebo without rehabilitation) in non-SCI cohorts results. Except for the two cases when the Tukey post hoc test was used for the PCR evaluation due to the reference of individual gene expression values to the untreated control without rehabilitation as well as for Max Speed test. Tukey test compares all possible pairs of means. In SCI cohorts results the Sidak’s multiple comparisons as recommended for pairwise comparisons of groups. The total amount of GAGs was assessed by using the one-way *ANOVA* test followed by Dunnett’s test multiple comparisons test. All the presented data in graphs were expressed as arithmetical means, with the standard error of the mean included. Significance as determined as followings: ns-not significant, *p < 0.05 **p < 0.01 ***p < 0.001 and ****p < 0.0001. Data were not assessed for normality. No test for outliers was conducted.

## RESULTS

### 4-MU decreases GAG synthesis and PNN in uninjured spinal cord

4-MU has been previously reported as a HA synthesis inhibitor (Nagy *et al*. 2015). HA is produced by the HAS enzymes from UDP-glucuronic acid (UDP-GlcA) and UDP-N-acetyl-glucosamine (UDP-GlcNAc). UDP-GlcA and UDP-GlcNAc are generated by UDP-glucuronosyltransferase which transfers an UDP-residue to N-acetylglucosamine and glucuronic acid (Bodevin-Authelet *et al*. 2005). 4-MU binds to glucuronic acid via the UGT (UDP-glucuronyl transferase). As a result, the concentration of UDP-GlcA decreases in the cytosol and HA synthesis is reduced (Nagy *et al*. 2015). While UDP-GlcA is a key substrate for HA synthesis, it is also the key substrate for CS, dermatan sulphate and heparan sulphate. Here, we first investigated the effect of 4-MU in the down-regulation of GAGs in the CNS and its effect on PNNs, with and without treadmill training which is a common rehabilitation. In addition, we tested the efficacy of a lower dose at 1.2 g/kg/day weight aiming to minimize the 4-MU consumption for potential adverse effects of long-term treatment for SCI (Dubisova *et al*. 2022).

Adult rats were fed with 4-MU at a dose of 1.2 g/kg/day daily for 8-weeks, and some of them were then subjected to 2 months of wash-out. We then extracted the GAGs from the dissected spinal cords and quantified the total amount for HA and GAGs using turbidity assay (Kwok *et al*. 2015) (Figure 2A). The results showed that 4-MU treated alone (0.09±0.81 mg/g; *n*=4; *p*=0.0086) and 4-MU plus daily treadmill training (0.17±0.16 mg/g; *n*=4; *p*=0.0113) have significantly reduced the level of GAGs compared to placebo group (2.02±0.70 mg/g; *n*=4). Rehabilitation did not affect the effectiveness of 4-MU in down-regulating GAG synthesis in treated animals. Interestingly, daily treadmill training also showed slightly, but non-significantly decrease in GAGs level in combination with placebo (1.06±0.16 mg/g; n=4; *p*=0.2555) suggesting that rehabilitation (or training) can independently reduce the level of GAGs. In wash-out group, the total amount of GAGs (0.93±0.40 mg/g; *n*=4; *p*=0.1734) recovered to a level similar to the rehabilitating group, suggesting a partial return of GAGs (Figure 2A). 4-MU and/or rehabilitative effect on GAGs level was test against placebo group.

**Figure 2.**
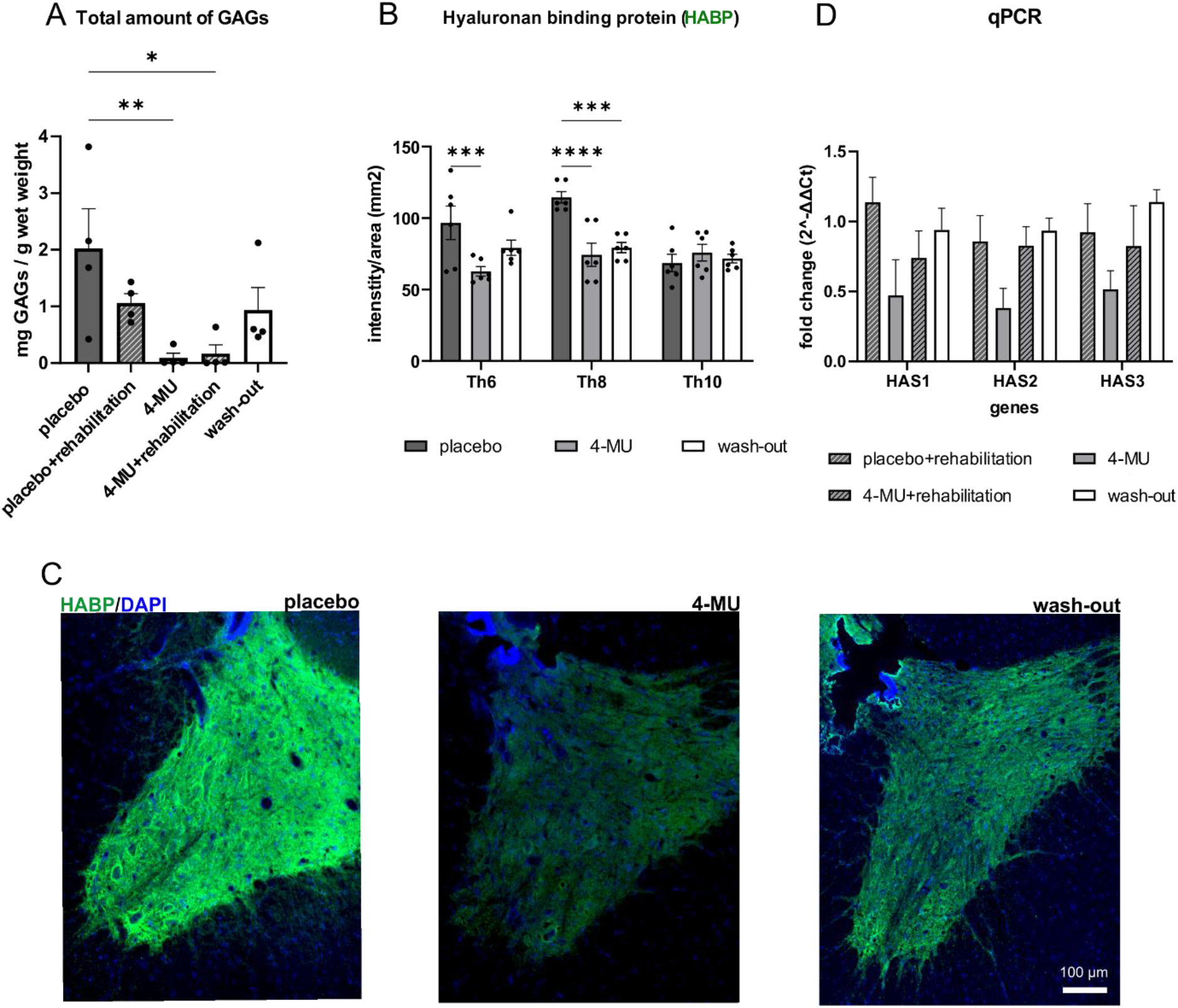
4-MU decreases hyaluronan (HA) and chondroitin sulphate proteoglycans (CSPGs) synthesis in non-SCI animals. (**A**) Bar graph showing total amount of glycosaminoglycans (GAGs) extracted from frozen spinal cords after 4-MU/placebo feeding and wash-out period. Values are plotted as mean ± SEM; * P < 0.05, by one-way ANOVA, Dunnett’s post-hoc test. (*n*= 4 animals per group). (**B**) Quantification of (C). Bar graph shows the mean intensity per area of grey matter together with the individual data points. Intensity was calculated using the HistoQuest software from TissueGnostics. Values are plotted as mean ± SEM; * *p* < 0.05, \*\**p* < 0.01 by two-way ANOVA, Dunnett’s post-hoc test. (*n*= 3 animals per group, 2 images per animal). (**C**) Representative fluorescent images showing different hyaluronan binding protein positive (HABP+) signal intensity in gray matter in placebo, 4-MU treated and wash-out group after 8 weeks of feeding and after 2 months of wash-out period. Scale bar: 100 μm. (**D**) Bar graph show the expression fold changes of the 2^-ΔΔCt values of the *hyaluronan synthase (HAS) 1, 2, 3* genes in comparison to the healthy and untreated animals. 2^-ΔΔCt values were determined by qRT-PCR. Values are plotted as mean ± SEM; *two-way ANOVA*, Tukey post-hoc test in all 4 groups. (*n*= 4 animals per group).

We also quantified the level of HA down-regulation using HABP staining (Figure 2, B and C). Histochemical staining was performed on sections at Th6 and Th10 and around the Th8 level which is the focus of subsequent experiments. In 4-MU treated group, the intensity of HABP was significantly decreased at Th8 level (74.32±12.81; *n*=3; *p*<0.0001) and at Th6 sections (68.68±4.20; *n*=3; *p*=0.0007) compared to placebo group (at Th8, 114.53±6.36, *n*=3 and at Th6, 114.57±10.97, *n*=3). There was no significant difference between 4-MU-treated and placebo groups in sections at Th10 level. After 2 months of wash-out, HA remained low at Th8 (79.37±5.57; *n*=3; *p*=0.0005) and at Th6 level (83.24±11.00; *n*=3; *p*=0.0962) compared to placebo group, with some trend of a return of HA production which did not reach the level of the placebo group (Figure 2, B and C).

Quantification of the mRNA expression of HASs in spinal cord samples using qPCR (Figure 2D) revealed no significant down-regulation of *HAS genes* expression in the 4-MU. However, clear trends were observed for *HAS genes* expression in 4-MU group when compared to all other groups. During the wash-out group the *HAS genes* expression reached the placebo combined with rehabilitation group level suggesting a recovery of the normal GAGs expression after wash-out period.

We next investigated the effect of 4-MU treatment (at a dose of 1.2 g/kg/day) on PNNs in the ventral horns by co-staining of WFA and ACAN. WFA is a widely used PNN marker and has been shown to specifically label the N-acetyl-D-galactosamine residue at terminal ends of chondroitin sulfate chains (Reichelt *et al*. 2019; Testa *et al*. 2019). ACAN is a major PNN component and has been reported to be superior in labelling PNN positive motoneurons in the spinal cord (Irvine and Kwok 2018). Spinal cord sections from all 5 groups (*n*=3 animals per group; 3 section per animal) were stained for WFA (Figure 3A, green arrows) and ACAN (Figure 3A, red arrows), the number of positive cells in the ventral horns up to the central canal was counted (Figure 3, B-D). Similar to the biochemical assays, 4-MU treatment and treadmill exercise independently reduced the total number of WFA-positive cells in spinal ventral horns (Figure 3, B-D). The combination of both induced a stronger down-regulation, however, this did not reach significance. In addition, we also observed the number of WFA and ACAN positive neurones returned to control levels after wash-out period. PNNs have been previously shown to enwrap α-motoneurons in the spinal cord. With the use of anti-ChAT antibody, we observed reduced PNNs around ChAT-positive neurones in thoracic uninjured spinal cords after 4-MU treatment (Figure 3E). This suggests that the current 4-MU dose at 1.2 g/kg/day or rehabilitation can effectively reduce PNNs in the uninjured spinal cord.

**Figure 3.**
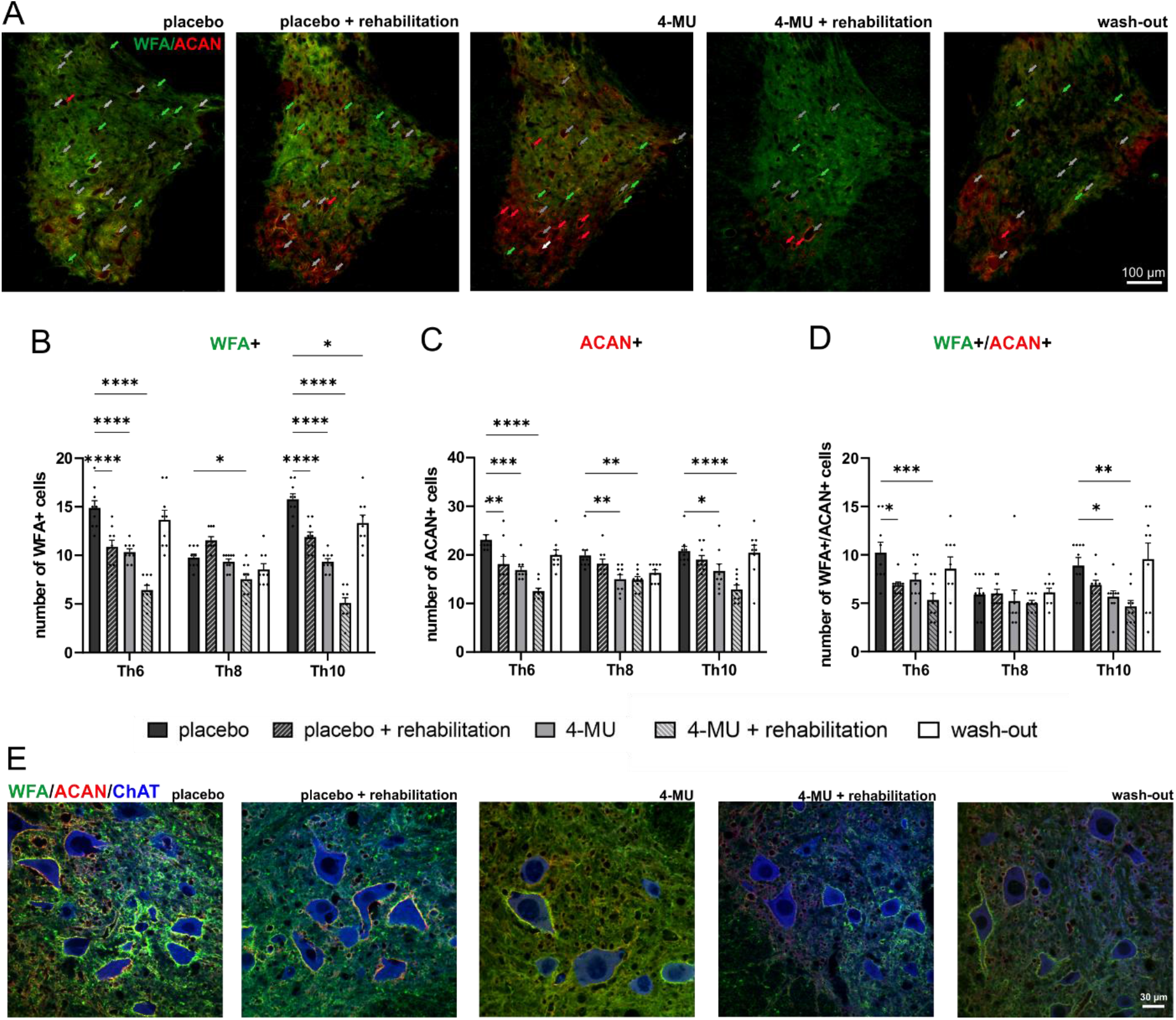
Down-regulation of perineuronal nets (PNNs) after 8 weeks of 4-methylumbelliferone (4-MU) feeding in non-spinal cord injury (SCI) animals, with or without daily treadmill training, and the re-appearance of perineuronal nets (PNNs) after 2 months of wash-out period. (A) Representative fluorescent images showing *Wisteria floribunda* (WFA) and aggrecan (ACAN) positive PNNs around cells in the ventral horns and their colocalization (WFA+/ACAN+) in thoracic spinal cord in uninjured animals in all 5 groups after 8 weeks of feeding and after 2 months of wash-out period. Green arrows indicated the WFA positive PNNs enwrapped cells; red arrows indicated ACAN positive PNNs enwrapped cells and gray arrows indicated cells where WFA/ACAN positive signal colocalizes. Scale bar: 100 μm. (B, C, D) Quantitative analysis of WFA positive or ACAN positive PNNs. Data showed mean ± SEM (*n*= 3 animals per groups, 3 sections per animal). \**p* < 0.05, \*\**p* < 0.01, \*\*\**p* < 0.001, **** *p* < 0.0001, two-way *ANOVA*, Dunnett’s multiple comparison. (E) Representative confocal images showing WFA positive and ACAN positive PNNs surrounding ChAT positive motoneurons in the thoracic rat spinal cord in placebo, 4-MU treated and wash-out group in ventral horn. Scale bar: 30 μm.

### 4-MU (1.2 g/kg/day) is insufficient to down-regulate the increasing production of chondroitin sulphates after SCI

After identifying that the 4-MU dose at 1.2 g/kg/day combined with rehabilitation can effectively reduce PNNs in the spinal cord, we next investigated if this dose is sufficient to reduce the increased expression of inhibitory ECM in injured spinal cord. Rats received 200 kdyn impact to the T8 spinal level which induced moderate SCI. Six weeks after the injury, animals were divided into two groups, groups were fed daily with 4-MU-containing or placebo chow for 8 weeks. In addition, both groups received task-specific rehabilitation for the 16 weeks concurrent to the oral 4-MU treatment (i.e. 8 weeks during 4-MU treatment and 8 weeks afterwards) in order to prime appropriate re-connection from the potentially heightened neuroplasticity. First, we evaluated the level of HA within the spinal cord (Figure 4) using HABP. There is a significant reduction of HA surrounding the lesion, rostrally (up to 5 mm) and caudally (1 and 5 mm) from the lesion in the 4-MU-treated group (40.78 ± 6.30) compared to placebo group (72.46 ± 6.46). We observed the trend of lower level of HA throughout the spinal cord after 4-MU treatment. The intensity remained decreased even after 8 weeks of wash-out period.

**Figure 4.**
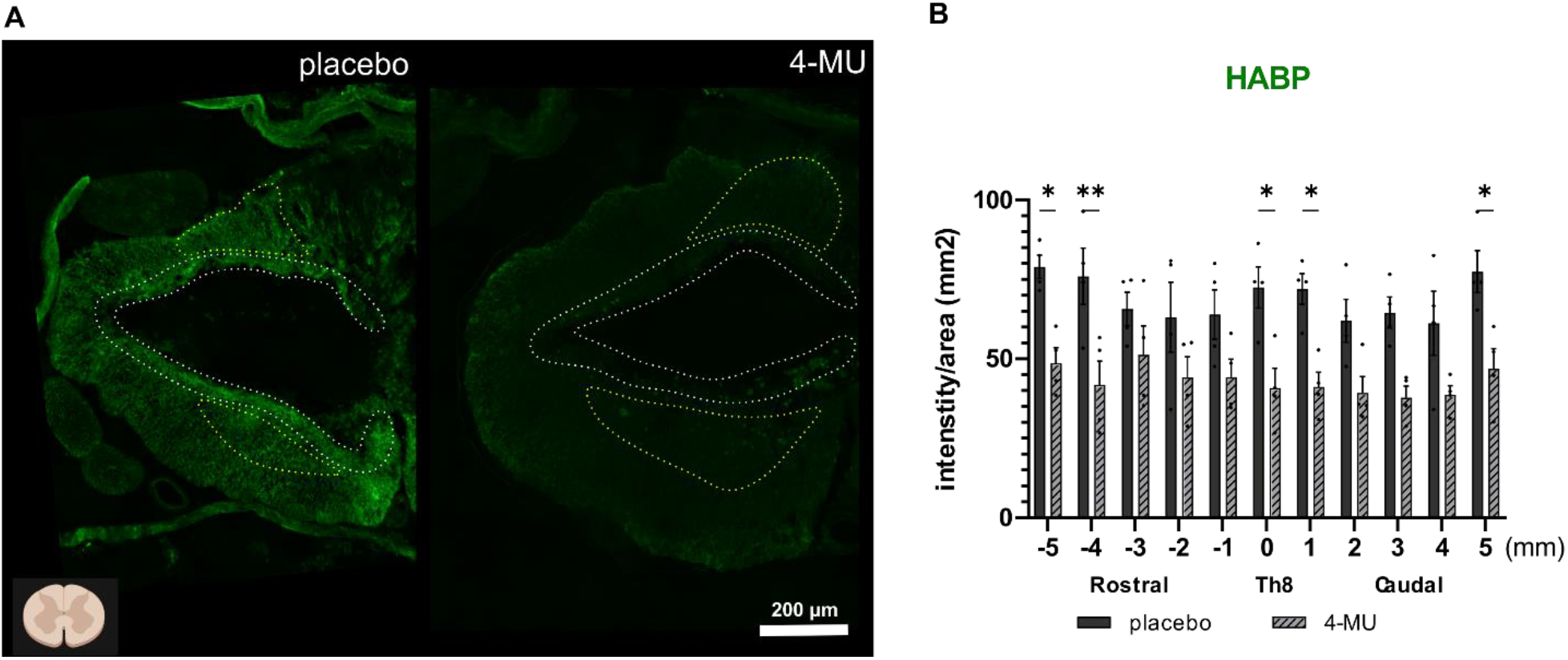
Hyaluronan binding protein (HABP) intensity remained decreased following chronic spinal cord injury 8 weeks of 4-MU treatment and 8 weeks wash-out period combined with daily rehabilitation. (**A**) Representative fluorescent images showing different HABP+ signal intensity in spinal cord injured (SCI) rats in placebo and 4-MU-treated groups after 8 weeks of feeding and after 8 weeks of wash-out period. Bar graph shows the intensity per sections throughout the spinal cord. White dotted lines delineating the border of the cavity. Yellow dotted lines depicting the spared gray matter. The area of the spared gray matter differs among animals due to injury variability and individual character of the animals. 3 animals were excluded based on BBB test 1 week after the injury (i.e. animals with less-then 8 score in BBB test were not included in the present study). Diagram in the bottom left shows the orientation of the cross sections; Created with BioRender.com. Scale bar: 200 μm; (**B**) Quantification of (A). Individual data together with their mean ± SEM were shown (*n*= 3 animals per group). * *p* < 0.05, ** *p* < 0.01, by two-way *ANOVA*,Sidak’s multiple comparisons test.

We then evaluated the level of PNNs and CSPGs using WFA and ACAN. Our results showed no significant difference between 4-MU-treated and placebo groups for 8 more weeks without any treatment but daily rehabilitation (Figure 5). These findings correspond with our data from uninjured animals where PNNs re-appeared after 2 months wash-out period (Figure 1 and 2). The strong up-regulation of CSPGs after SCI has rendered the dose of 1.2 g/kg/day 4-MU to be insufficient to supress their production in injured spinal cord.

**Figure 5.**
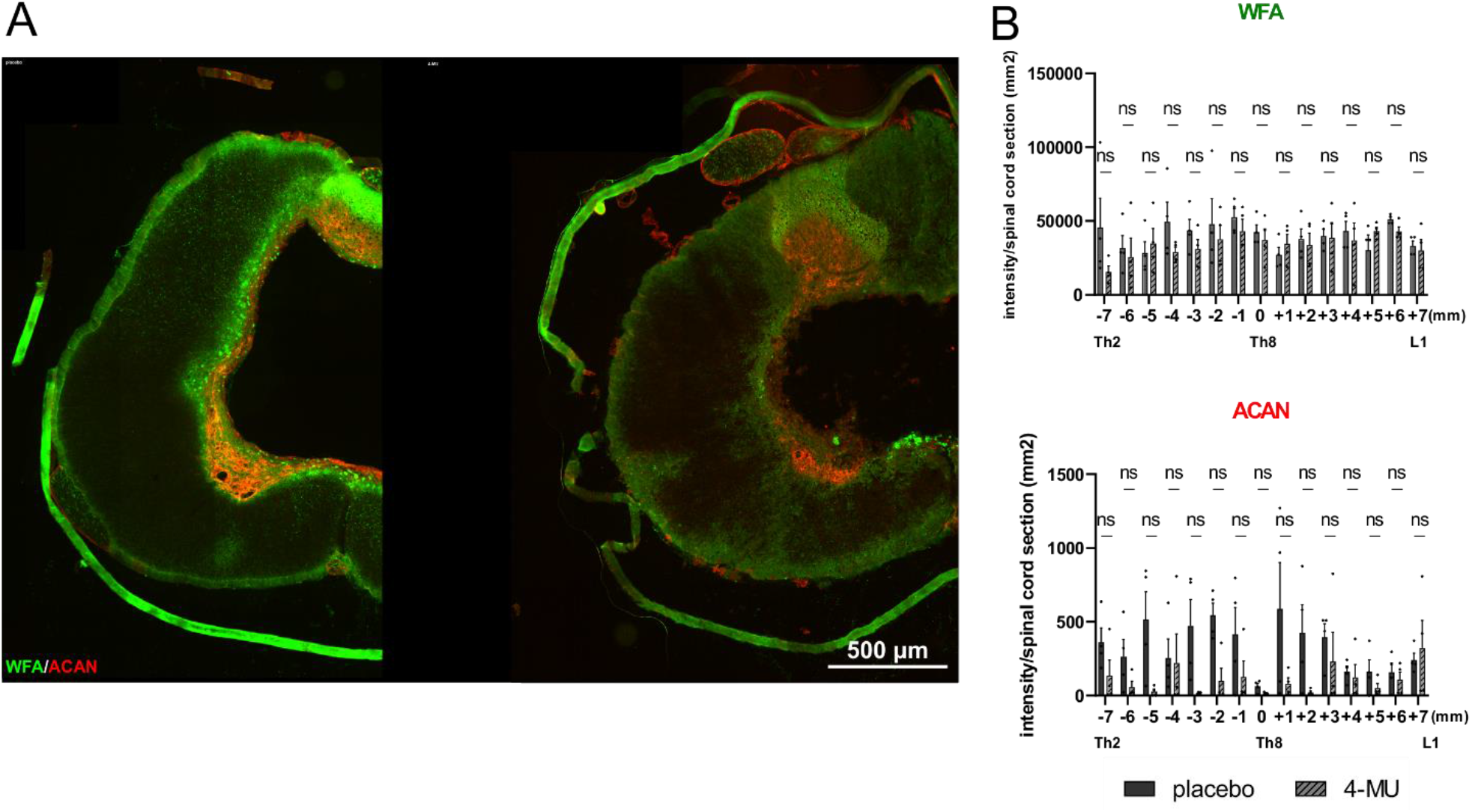
Immunofluorescence double-staining of *Wisteria floribunda* agglutinin (WFA) and aggrecan (ACAN) suggested that 4-MU at current dose 1.2 g/kg/day is not sufficient to down-regulate the increasing production of chondroitin sulphates after spinal cord injury (SCI). (**A**) Representative fluorescent images showing WFA+ (in green) and ACAN+ (in red) area around the centre of the lesion. Scale bar: 500 μm; (**B, C**) The quantitative analysis of WFA or ACAN intensity. Bar graphs show the intensity per sections throughout the spinal cord with the lesion centre marked as level 0. Individual data points and their mean ± SEM were shown (*n*= 4 animals per group). ns by two-way *ANOVA*,Sidak’s multiple comparisons test.

### 4-MU reduces glial scar in chronic spinal cord injury

HA is produced by both neurons and glia in the CNS, and is up-regulated by astrocytes in neuroinflammation (Asher *et al*. 1991; Struve *et al*. 2005; Back *et al*. 2005). The astrocyte activation and the formation of glial scar aim to limit spreading of neuroinflammation from the injury epicentre to the surrounding uninjured tissue (Wang *et al*. 2018). However, the glial scar creates a mechanical and biochemical barrier preventing spontaneous regeneration (Bradbury and Burnside 2019). It has been previously suggested that 4-MU reduces astrogliosis in the brain parenchyma in the mouse model of the experimental autoimmune encephalomyelitis (Kuipers *et al*. 2016). In view of our observation in HA down-regulation around the lesion cavity, we next focused on the effect of 4-MU to the glial scar after SCI. Quantitative analysis of GFAP-positive area was performed to assess the glial scar surrounding the lesion cavity on cross sections (Figure 6). A significant decrease in astrogliosis was observed in 4-MU treated group at a dose of 1.2 g/kg/day compared to placebo group. The average peak in the centre of the lesion in 4-MU groups was 1.49 ±0.51% (*n*=4) and 6.1 ±1.94% (*n*=4) in placebo groups (Figure 6C).

**Figure 6.**
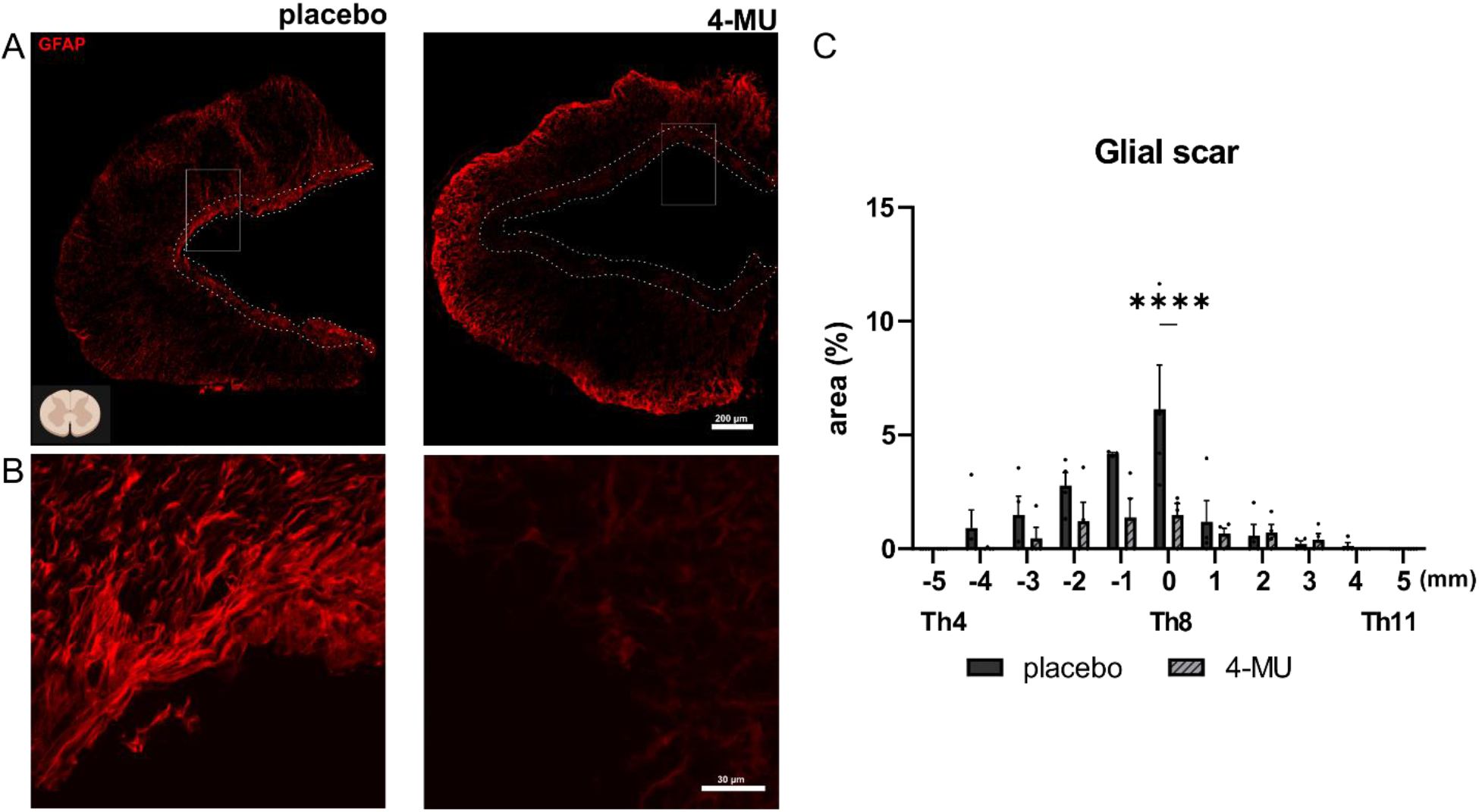
4-MU treatment reduced glial scar area surrounding the lesion site. (**A**) Representative fluorescent images showing lesion epicentre stained for glial fibrillary acidic protein (GFAP) in 4-MU treated and placebo group with chronic spinal cord injury. Dotted lines show the area border of the lesion cavity in 4-MU treated group and GFAP+ area in placebo group. Scale bar: 200 μm; Diagram of uninjured spinal cord at top left showing the direction of the cross section in (A), Created with BioRender.com; (**B**) Magnified images showing structural change of the glial scar tissue after 4-MU treatment compared to untreated animals. Scale bar: 30 μm; (**C**) Bar graph showing area of the glial scar around the central cavity performed in the GFAP stained histochemical images using ImageJ software. Values are plotted as mean ± SEM; **** *p* < 0.0001 by two-way ANOVA, Sidak post-hoc test. (*n*= 4 animals per group).

### Daily 4-MU at a dose of 1.2 g/kg/day promotes plasticity of serotonergic fibres in chronic spinal cord injury

4-MU has previously been shown to down-regulate PNNs in the brain at a dose of 2.4 g/kg/day (Dubisova et al., 2022). The results in figures 4 and 5 showed that while lower dose of 1.2 g/kg/day is able to reduce HA, the level of CSs remains similar to untreated injured controls after SCI (Figure 5). Previous work has shown that HA is involved in the sprouting of serotonergic fiber and thus enhance neuroplasticity (Barzilay *et al*. 2016). It has been previously suggested that 5-HT2C receptors are strongly involved in synaptic plasticity by initiating the phosphoinositol second messenger cascade through inositol triphosphate and diacylglycerol (Lesch and Waider, 2012). Here, we investigated the effect of 1.2 g/kg/day 4-MU treatment and its subsequent HA down-regulation on the expression of 5-HT2C receptors after SCI. Spinal cord sections were stained, rostrally and caudally from the Th8 lesion site, for 5-HT2C in the ventral horns in both treated and control groups (Figure 7A). After 8 weeks of 4-MU treatment and 2 months-long wash-out period plus rehabilitation, there is a significant increase of 5HT2C receptor immunoreactivity in the ventral horns above and below the lesion in the 4-MU-treated (13.85±0.86) when compared to the placebo group (10.19 ± 0.49) (Figure 7B). The results suggest that following chronic spinal cord injury, rehabilitation and 8 weeks wash-out period, 4-MU treatment and its down-regulation in HA promote long-lasting synaptic plasticity.

**Figure 7.**
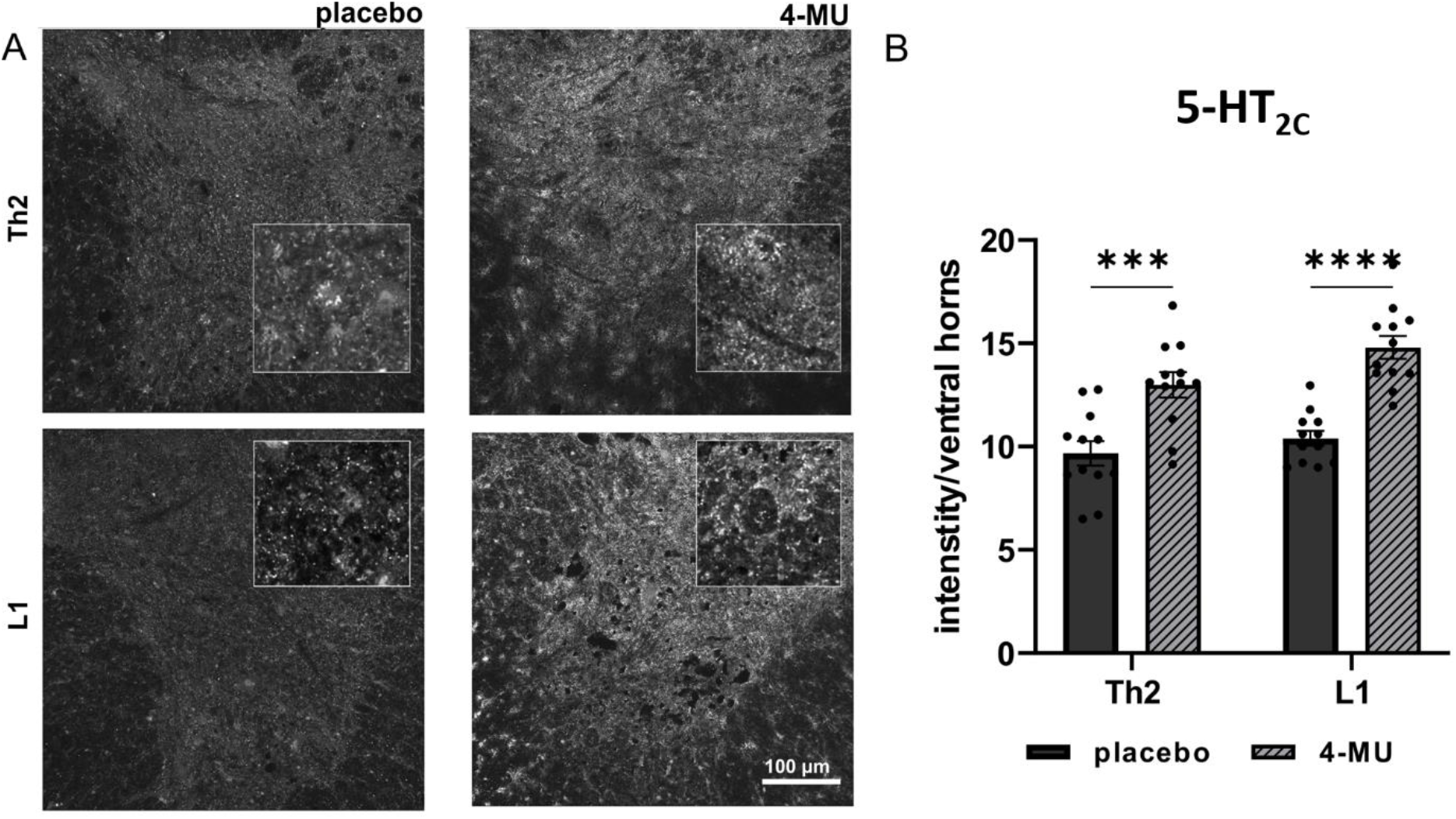
4-MU treatment promotes serotonergic fibre plasticity after chronic spinal cord injury (SCI). (**A**) Representative confocal images showing ventral horn below and above lesion stained for 5-hydroxytryptamine 2C receptor (5-HT_2C_) in 4-MU treated and placebo group in chronic stage of SCI. Insets show the magnified views of the staining. Scale bar: 100 μm; (**B**) Bar graph showing intensity measurement per area (in pixels) of ventral horns using ImageJ software. Values are plotted as mean ± SEM (*n*= 4 animals per group; 3 sections per animal); *** *p* < 0.001, \*\*\*\**p* < 0.0001 by two-way *ANOVA*, Sidak’s multiple comparisons test.

### 4-MU at a dose of 1.2 g/kg/day has no effect on synaptic density

To investigate whether there is a difference in overall synaptic density (Figure 8), we stained 2 levels above and below the lesion for the presynaptic marker synapsin and we measured the level of synaptic contacts within ventral horns. 4-MU treated animals showed a trend in increased synaptic density in the area above lesion, but with no significant difference (Figure 8B). There is no change in synaptic density measurement below lesion, probably due to the lack of CS reduction as observed in Figure 5.

**Figure 8.**
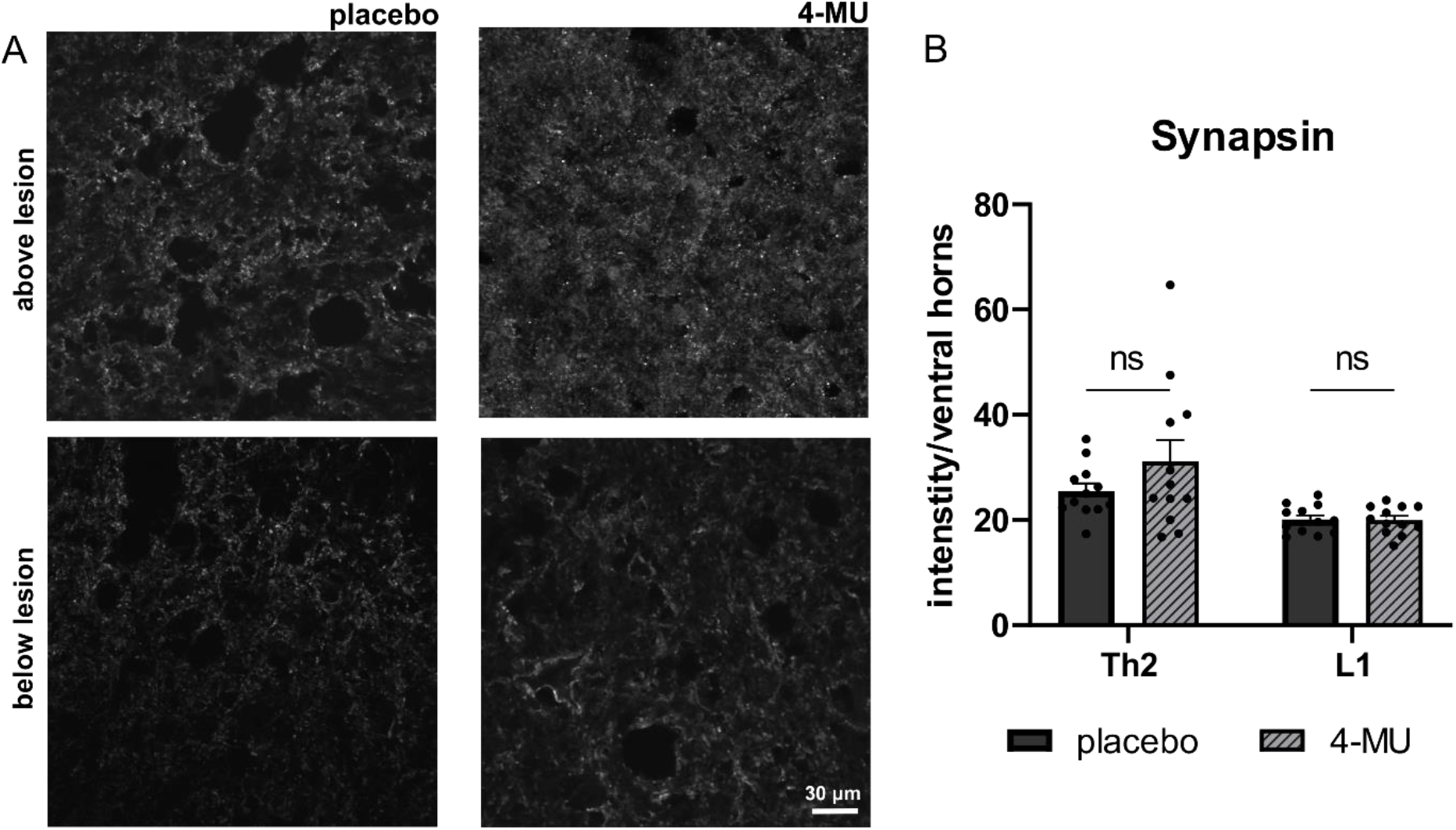
4-MU treatment does not affect synapse number. (**A**) Representative confocal detailed images showing ventral horn above and below lesion stained for synapsin in 4-MU treated and placebo group in chronic stage of spinal cord injury. Scale bar: 30 μm; (**B**) Bar graph showing intensity measurement per area (in pixels) of ventral horns using ImageJ software. Values are plotted as mean ± SEM (*n*= 4 animals per group; 3 sections per animal); ns > 0.05 by two-way *ANOVA*, Sidak’s multiple comparisons test.

### 4-MU at a dose of 1.2 g/kg/day is not sufficient to enhance inherent functional recovery in chronic stage of spinal cord injury

On account of biochemical results showing that 4-MU abolishes plasticity-limiting perineuronal nets, we tested if 4-MU combined with daily rehabilitation induce axonal sprouting leading to functional recovery in chronic stage of SCI. To test this, the whole battery of behavioral test was chosen, including weekly BBB test, max speed test and ladder rung walking test combined with daily rehabilitation on treadmill assessing their locomotor abilities. Two sensory tests, mechanical pressure test (Von Frey test) and thermal test (Plantar test) were chosen to assess changes in thermal and mechanical sensation. These two tests were performed just trice - before feeding with 4-MU-containing / non-containing pellets, at the end of feeding period, and at the end of the whole experiment. In our results, there were no significant differences nor indication for a trend of improvement at any time point suggesting 4-MU-mediated recovery (Figure 9).

**Figure 9.**
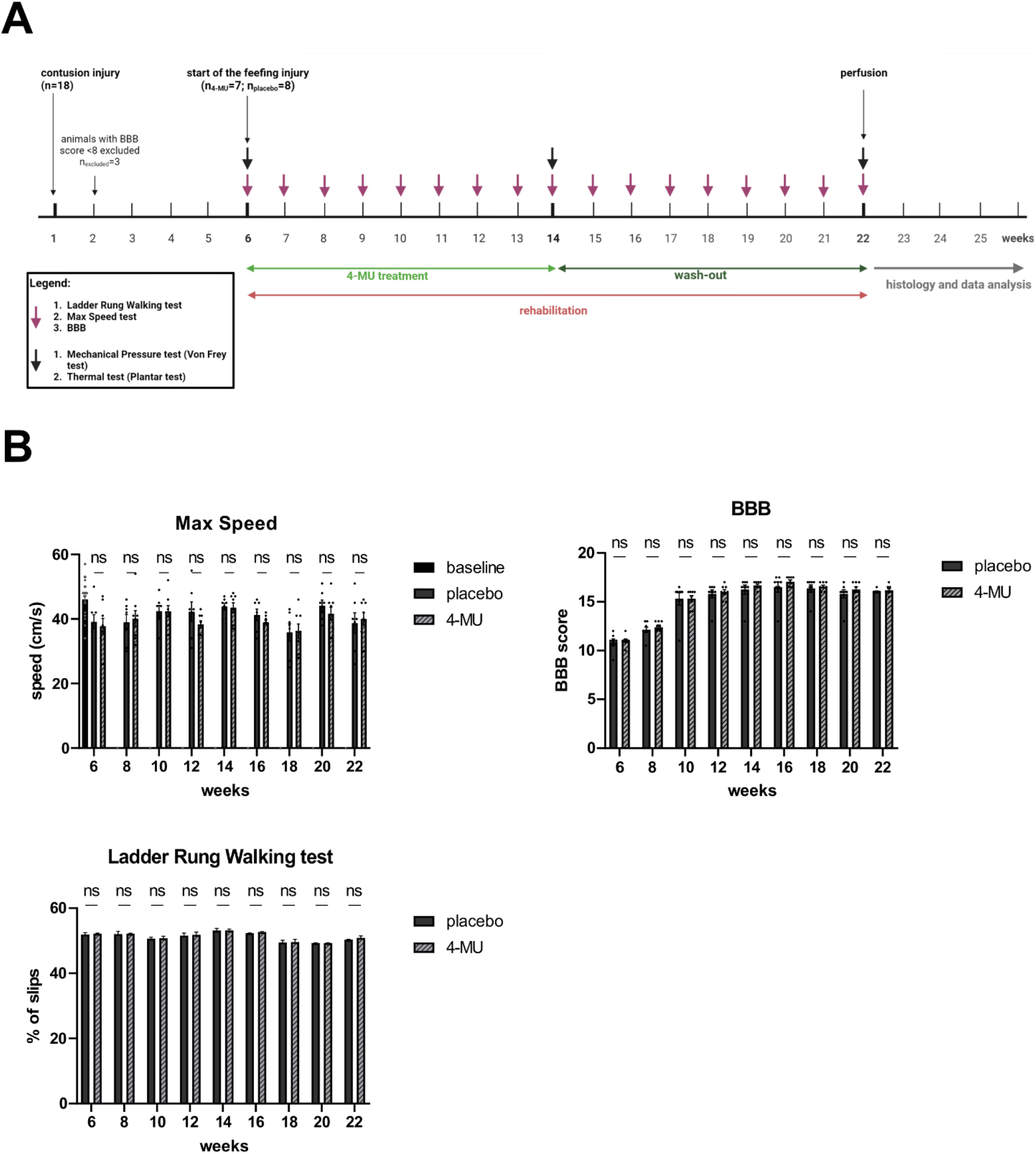
4-MU treatment at a current dose does not lead to functional recovery after chronic SCI. Animals were assessed weekly for their ability to reach the highest possible speed on the treadmill (Max Speed), to locomote in the Basso, Beattie, Bresnahan (BBB) open-field test, and to walk skilfully and place paws correctly (Ladder rung walking test). **(A)**shows the timeline when the tests were performed; created with BioRender.com. **(B)**bar graphs display spontaneous recovery in both groups after first two weeks of daily rehabilitation in BBB open-field test that plateaued during the week 10 in both groups. There was no 4-MU-mediated enhancement of this spontaneous recovery. Values are plotted as mean ± SEM (*n*= 7 animals for placebo group and *n*= 8 animals for 4-MU group); ns > 0.05 by two-way *ANOVA*, Sidak’s multiple comparisons test (BBB, Ladder Rung Walking test); ns > 0.05 by two-way *ANOVA*, Tukey’s multiple comparisons test (Max Speed).

## DISCUSSION

We investigated whether a dose (1.2 g/kg/day) of 4-MU was sufficient to decrease PNNs in ventral horns, and to promote sprouting and functional recovery in chronic SCI. Previous study in mice using 2.4 g/kg/day has led to an enhancement neuroplasticity for memory acquisition through PNN down-regulation. The dose of 2.4 g/kg/day is equivalent to a quarter of the LD50. We thus tested in this study if halfing this dose at 1.2 g/kg/day will be able to down-regulate PNNs for potential functional recovery after SCI. We observed that a 1.2 g/kg/day dose of orally-administered 4-MU is sufficient to reduce PNNs and HA levels in uninjured animals, but seems not to be sufficient enough to suppress strong CSPGs upregulation after SCI, and thus not allowing functional recovery.

In recent years, many studies have shown strategies focusing on regeneration after SCI, often based on targeting PNNs and manipulating the glial scar to attenuate inhibitory properties of its environment. Current strategies range from proteolytic ECM manipulation to targeting the specific ECM components through the synthesis of inhibitory ECM molecules after SCI (Burnside and Bradbury 2014). One of the most studied approaches has been an enzymatic ECM modification using ChABC. ChABC degrades CS chains into disaccharides and removes CSPG inhibition in the glial scar as well as removes the PNNs as plasticity brake, to benefit both acute and chronic SCI conditions (Wang *et al*. 2011). Animals with SCI show better recovery both anatomically and functionally after ChABC treatment (Bradbury *et al*. 2002; García-Alías *et al*. 2009; Wang *et al*. 2011). Functional recovery after SCI is further boosted when plasticity restoration is combined with rehabilitation (García-Alías *et al*. 2009; Wang *et al*. 2011). However, ChABC application has several disadvantages. The major disadvantage of using this bacterial enzyme is its thermal instability and short half-life, which requires multiple or continuous intrathecal administrations (Nori *et al*. 2018), potential immune response of the body and difficult dosing. Apart from ChABC, Keough *et al*. (2016) were testing a subset of 245 drugs known for their CNS penetrating capacity and oral bioavailability. None of these 245 compounds showed sufficient ability to overcome CSPG inhibition from oligodendrocyte precursor cells.

4-MU, the derivative of coumarin, used for biliary stasis therapy, is a HA inhibitor (Nagy *et al*. 2015). It has been previously shown to reduce the synthesis of HA by decreasing the production of UDP-GlcA, a key monosaccharide substrate for the synthesis of HA (Weigel 2015), and the expression of HASs. As UDP-GlcA is also a substrate for CS production, we thus investigate if 4-MU administration would decrease the synthesis of both HA and CS, facilitating neuroplasticity. Indeed, we have observed the down-regulation of HA and CS, both anatomically using histochemistry and biochemically through GAG quantification. Our data suggest that 4-MU combined with daily training suppresses the synthesis of GAG. With the use of PNN markers including WFA and ACAN, 4-MU administration led to PNNs removal in ventral horns. PNNs re-appeared after 2 months of wash-out period. The reason of why PNNs reappeared while the HA level remained low after 2 months of wash-out is likely because CS synthesis is less sensitive to the UDP-GlcA deficiency. CS is synthesized in the Golgi apparatus where UDP-GlcA sugars are being transported to the Golgi lumen with high affinity, while HA is synthesized directly at the cytoplasmic membrane (Nagy *et al*.2015).

We used a thoracic contusion injury that spares some axons around a central cavity and ablates dorsal corticospinal tracts (CSTs) critical for motor control in humans, thus mimics the type of closed SCI most commonly seen in humans (Basso 2000). The 4-MU treatment started in the chronic stage, *i.e*. 6 weeks after the injury (Kjell and Olson 2016). At this stage, glial scar is well-established, CSPGs are upregulated, and the acute immune response had already subsided (Stichel and Müller 1994; Hu *et al*. 2010). 4-MU treatment was accompanied by daily rehabilitation on treadmill to consolidate appropriate synaptic connections and prune away others (Oudega *et al*. 2012). After the 8 weeks of treatment and rehabilitation, rehabilitation was continued for another 8 weeks for the wash-out. This wash-out period gives time for PNNs to re-form and stabilize *de novo* synapses and consolidate anatomical plasticity (García-Alías *et al*. 2009; Wang *et al*.2011; Al’joboori *et al*. 2020), while the continuing rehabilitation prunes random connections, supporting appropriate connections and removing inappropriate ones (Fawcett and Curt 2009; Kanagal and Muir 2009). Oral administration of 1.2 g/kg/day 4-MU robustly decreased the glial scar surrounding the cavity that persisted throughout the wash-out period.

To evaluate the potential of 4-MU to remove the plasticity brake formed by PNNs, we analysed the intensity of 5-HT signal and observed increased 5-HT sprouting above and below the lesion. However, this sprouting did not lead to any significant difference in synapsin immunoreactivity within the ventral horns, above and below the lesion, between the 4-MU-treated and placebo animals. Assessment of functional recovery using the BBB test, maximum speed test and ladder rung walking test yielded no significant differences between the 4-MU and placebo treated groups (data not shown), even with the continuous rehabilitation for next 2 months. As 4-MU-mediated PNNs ablation has been already demonstrated in previous work focused on mouse hippocampus where it enhances memory in ageing mice (Dubisova *et al*. 2022) we reasoned that the lack of functional recovery after SCI in this study is due to the lower dose of 4-MU administered (2.4 g/kg/day *versus* 1.2 g/kg/day). A higher dose of 4-MU combined with rehabilitation should be tried for its effect on recovery after SCI.

The promising results we observed in 5-HT2C staining agree with published data. Previous studies have already demonstrated that severe SCI causes paralysis not only mediated by a loss of direct muscle innervation from spinal motoneurons, but also by loss in supraspinal tracts involved in voluntary initiation of movements and by loss in descending tracts providing motoneurons neuromodulators such as serotonin (5-HT) (Jacobs *et al*. 2002; Jordan *et al*. 2008). 5-HT stimulates spinal interneurons and motoneurons, thus allows appropriate muscle contraction (Jacobs *et al*. 2002; Perrier and Delgado-Lezama 2005). In the acute SCI, spinal motoneurons and interneurons lack 5-HT (Perrier and Hounsgaard 2003; Harvey *et al*. 2006) resulting in paralysis and spinal shock (Bennett *et al*. 1999; Bennett *et al*. 2004). It has been showed that motoneurons are able to adapt to the lack of 5-HT due to constitutive action of 5-HT2C receptors. Therefore, in chronic SCI, motoneurons partially recover their excitability, in spite of the continued absence of 5-HT (Button *et al*. 2008). It was also suggested that this compensatory mechanism needs activity-dependent tuning to contribute to locomotor activity after SCI (Murray *et al*. 2010).

In order to determine if there is any compensatory mechanism for the loss of HA synthesis, we have evaluated the changes in mRNA expression for genes related to the HA synthesis (*has1, has2, has3*) (Nagy *et al*. 2015). We showed a significant down-regulation in *has1* and clear trend in *has2* mRNA levels. The *has* genes expression after 1 mM 4-MU treatment, studied on human aortic smooth muscle cells showed reduction mainly in *has1* and *has2* transcripts after 4-MU treatment (Vigetti *et al*. 2009). To determine the biochemical effect of 4-MU at 1.2 g/kg/day on the HA synthesis in uninjured animals, we used histochemical staining for recombinant HABP derived from human versican G1 domain which binds specifically to hyaluronan and does not bind to other glycosaminoglycans (Amsbio; data sheet). We observed that after 2 months wash-out period, the level of HA remains decreased compared to healthy animals. This may pose a question to whether the HA down-regulation after 2 months of wash-out period may induce adverse effects after treatment.

Clinical trials with 4-MU (drug approved in Europe and Asia as hymecromone) showed excellent safety parameters during long-term administration of approved doses – 1.2 – 3.6 g/day (Abate *et al*. 2001; Camarri and Marchettini 1988; Krawzak *et al*. 1995; Quaranta *et al*. 1984; Trabucchi *et al*. 1986; Walter and Seidel 1979; Hoffmann *et al*. 2005). The longest reported duration of oral 4-MU administration was for 3 months in human patients (Trabucchi *et al*. 1986) which is much shorter time according to correlation between age of laboratory rats and humans (Sengupta 2013) and it might be taken into consideration while explaining these results as the safety profile of long-term 4-MU treatment is not fully established yet (Nagy *et al*. 2015).

In conclusion, this study demonstrates that oral administration of 1.2 g/kg/day of 4-MU reduce the total amount of GAGs, reduce glial scar and increase 5-HT sprouting in chronic stage of SCI. However, these structural changes do not result in increased synaptic connections and do not lead to functional recovery even after intensive rehabilitation. A higher dose of 4-MU is likely to be necessary to induce the functional recovery supported by synaptic plasticity.

## List of abbreviations

4-MU: 4-methylumbelliferone
5-HT: 5-hydroxytryptamine
ACAN: aggrecan
ChAT: choline acetyltransferase
ChABC: chondroitinase ABC
CNS: central nervous system
CS-GAGs: chondroitin sulfate glycosaminoglycans
CSPGs: chondroitin sulfate proteoglycans
HA: hyaluronan
HABP: hyaluronan binding protein
HASs: hyaluronan synthases
UDP-GlcA: UDP-glucuronic acid
UGT: UDP-glucuronyl transferase
PNNs: perineuronal nets
SCI: spinal cord injury
WFA: Wisteria floribunda agglutinin

## Acknowledgements

Supported by: Center of Reconstruction Neuroscience – NEURORECON CZ.02.1.01/0.0/0.0/15_003/0000419 (to LMU, PV and JCFK) and Czech Science Agency 19-10365S (to PV and JCFK). Wings for Life (WFL-UK-008-15) and Medical Research Council UK (Confidence in concept MC-PC-16050 and Project grant MR/S011110/1) to JCFK.

## Author contributions

SK, LMU, PJ and JCFK conceived and designed the study, and obtained funding. KS, MC, ZS, NMV and LMU performed the experiments, analysed and interpreted the data. All authors are involved in writing and proof-reading the manuscript.

